# CRISPR/Cas9-mediated disruption of *CjACOS5* confers no-pollen formation on sugi trees (*Cryptomeria japonica* D. Don)

**DOI:** 10.1101/2023.01.16.521755

**Authors:** Mitsuru Nishiguchi, Norihiro Futamura, Masaki Endo, Masafumi Mikami, Seiichi Toki, Shin-Ichiro Katahata, Yasunori Ohmiya, Ken-ichi Konagaya, Yoshihiko Nanasato, Toru Taniguchi, Tsuyoshi Emilio Maruyama

## Abstract

Sugi (*Cryptomeria japonica* D. Don) is an economically important coniferous tree in Japan. However, abundant sugi pollen grains are dispersed and transported by the wind each spring and cause a severe pollen allergy syndrome (Japanese cedar pollinosis). The use of pollen-free sugi that cannot produce pollen has been thought as a countermeasure to Japanese cedar pollinosis. The sugi *CjACOS5* gene is an ortholog of *Arabidopsis ACOS5* and rice *OsACOS12*, which encode an acyl-CoA synthetase that is involved in the synthesis of sporopollenin in pollen walls. To generate pollen-free sugi, we mutated *CjACOS5* using the CRISPR/Cas9 system. As a result of sugi transformation mediated by *Agrobacterium tumefaciens* harboring the *CjACOS5-targeted* CRISPR/Cas9 vector, 1 bp-deleted homo biallelic mutant lines were obtained. Chimeric mutant lines harboring both mutant and wild-type *CjACOS5* genes were also generated. The homo biallelic mutant lines had no-pollen in male strobili, whereas chimeric mutant lines had male strobili with or without pollen grains. Our results suggest that *CjACOS5* is essential for the production of pollen in sugi and that its disruption is useful for the generation of pollen-free sugi. In addition to conventional transgenic technology, genome editing technology, including CRISPR/Cas9, can confer new traits on sugi.

## Introduction

Sugi (Japanese cedar, *Cryptomeria japonica* D. Don) is a conifer species in the family Cupressaceae in gymnosperms^1^. It is one of the important domestic trees in Japan for forestry, industry, and the economy. Artificially sugi-planted areas had reached 4.44 million ha in 2017, covering about 12% of the land area of Japan^2^. The production of sugi roundwood was 12.28 million m^3^, which represented the largest volume (57% of the total volume) among the domestic tree species in 2017^3^. Sugi wood is widely used as a structural, construction, and packaging material, as well as for flooring, ceiling boards, barrels, chopsticks, etc.

Conversely, sugi pollen allergy syndrome (Japanese cedar pollinosis) is a serious disease in Japan^4^. Sugi produces male strobili and female strobili, similar to other monoecious conifer trees. Sugi pollen is dispersed from male strobili in early spring and is carried far away by the wind. The pollen grain contains multiple allergen proteins: Cry J 1 (pectate lyase), Cry J 2 (polygalacturonase), Cry J 3 (thaumatin-like protein), CJP-4 (class IV chitinase), CJP-6 and CJP-8 (isoflavone reductase-like lipid transfer proteins), CPA9 (subtilisin-like serine protease), and CPA63 (aspartic protease)^4^. The pollen grains enter the human body and adhere to the nasal and ocular mucosa. They are burst by water absorption, release cytoplasmic components, including the allergens, and induce type II immunity. Japanese cedar pollinosis was first reported in 1964 in Nikko, Japan^5^. Thereafter, several nationwide surveys showed that the number of patients with Japanese cedar pollinosis had been increasing: the estimated prevalence was 11.7% in 1998^6^, 13.1% in 2001^7^, 26.5% in 2008^8^, and 38.8% in 2019^9^.

The reduction of the amount of sugi pollen has been thought as a countermeasure to Japanese cedar pollinosis. For this purpose, wild varieties with fewer male strobili and wild male-sterile mutants without pollen grains have been used for breeding. Male-sterile mutants were discovered in many plant species and the genetic patterns of male sterility are divided into genic male sterility (GMS) and cytoplasmic male sterility^10,11^. GMS is also called nuclear male sterility or Mendelian sterility because it depends exclusively on the nuclear genome and exhibits Mendelian inheritance. The development of genetic analyses has served to identify several key genes involved in GMS. For example, a transposon insertion mutant of the *Arabidopsis thaliana* acyl-CoA synthetase 5 (*ACOS5*) gene does not produce mature pollen in anthers^12^. The recombinant ACOS5 protein catalyzes oleic acid CoA-ester formation *in vitro*, and thus probably participates in the biosynthesis of sporopollenin, which is a constituent of the exine of pollen grains. Rice (*Oryza sativa*) carries the *OsACOS12* gene, which is an ortholog of *ACOS5*, and the *osacos12* mutant also shows a phenotype of absence of mature pollen^13,14^.

A male-sterile sugi without pollen was first identified in Japan in 1992^15,16^; subsequently, various male-sterile mutant trees were discovered and analyzed^17–19^. To identify the genes that are responsible for male sterility in sugi, genetic marker analyses have been performed, leading to the identification of four recessive male sterility-linked independent loci (*ms1, ms2, ms3* and *ms4*)^20–24^. Recently, one gene (*MS1*) at the *ms1* locus was identified as a causative gene for male sterility^25^. *MS1* encodes a lipid transfer protein and is expressed in male strobili, but its biochemical properties or physiological roles remain unknown. The appearance rate of pollen-free sugi trees is approximately 0.02% in a seed orchard in Japan^26^, and these trees are extremely rare in the field. In addition, the discovered mutant trees should be crossed to sugi plus trees with excellent properties for the generation of pollen-free plus trees, because most of the original male-sterile mutant trees exhibit slow growth or curved trunks. Therefore, the development of male-sterilization technology is valuable and needed for the generation of pollen-free sugi plus trees.

In this study, we aimed to generate pollen-free sugi trees using the CRISPR/Cas9 system. Genome editing technology including the CRISPR/Cas9 system has been widely utilized in plants and animals for the elucidation of gene function and for the breeding of new varieties. Because of the simplicity of its experimental procedures, the CRISPR/Cas9 system has been applied not only to herbaceous plants and crops, but also to woody plants such as sweet orange trees^27^, apple trees^28^, grape vines^29^, poplar trees^30,31^, sugi trees^32^, radiata pine^33^, and white spruce^34^. Although the use of genetically modified organisms (GMOs) including transgenic trees have been regulated by law in Japan, the products by the genome editing technology are less restricted by law, if they have no DNA from other species. Recently, γ-aminobutyric acid (GABA)-enriched tomato which had been generated by CRISPR/Cas9 started to be commercially sold without the regulation against GMOs^35^. The genome editing technology has the potential to put the modified and improved forest trees to practical use in future. Here we report the genome editing of sugi by the CRISPR/Cas9 system, to mutate the *CjACOS5* gene, which is an ortholog of *Arabidopsis ACOS5* and rice *OsACOS12*. The regenerated sugi trees with mutated *CjACOS5* genes flowered but did not produce pollen grains in male strobili. We demonstrated that the CRISPR/Cas9 system enables the generation of pollen-free sugi trees artificially, even though we had not been able to achieve this in the past.

## Results

### Isolation of the *CjACOS5* genes from sugi

To induce male sterility in sugi, we searched for the target genes that are disrupted using the CRISPR/Cas9 system. We noticed that the *A. thaliana acos5* mutant and the rice *osacos12* mutant have been reported to exhibit male sterility and not to produce pollen^12,13^. Both the *ACOS5* (TAIR ID: AT1g62940) and *OsACOS12* (MSU ID: LOC_Os04g24530, RAP-DB ID: Os04g0310800) genes encode an acyl-CoA synthetase that is involved in sporopollenin synthesis during pollen formation. We searched for *ACOS5*-homologous genes in full-length cDNA libraries of sugi male strobili^36^, and identified one cDNA, CMFL003_A04 (DDBJ accession number: FX341539). The NCBI blastn program showed that CMFL003_A04 was homologous to *Phoenix dactylifera* 4-coumarate-CoA ligase-like 1 (GenBank accession: XM_008810877, e-value: 1e-81), *Amborella trichopoda* 4-coumarate-CoA ligase-like 1 (XM_006850938, 7e-55), *A. thaliana ACOS5* (AY250836, 3e-53), *Capsella rubella* 4-coumarate-CoA ligase-like 1 (XM_006301215, 4e-52), etc.

Furthermore, we cloned cDNAs homologous to CMFL003_A04 from male strobili of sugi #13-8-2 line and sequenced. As a result, we identified two cDNAs, *CjACOS5a* (accession number: LC726337) and *CjACOS5b* (LC726338). The predicted protein encoded by *CjACOS5a* and *CjACOS5b* comprises 557 amino acid residues and their 551 residues (98.9%) are equal to each other (Fig. 1). The CjACOS5a protein has 61.9% identity to ACOS5 and 59.8% identity to OsACOS12. The CjACOS5 proteins are composed of two domains; one is an AMP-dependent synthetase/ligase domain (InterPro ID: IPR000873), from Glu41 to Tyr454, and the other domain is a C-terminal domain in AMP-binding enzymes (IPR025110), from Glu463 to Lys538 (Fig. 1). Both domains also exist in ACOS5 and OsACOS12. In the AMP-dependent synthetase/ligase domain, the CjACOS5 proteins have a conserved AMP-binding site (IPR020845), from Leu197 to Lys208 (LPYSSGTTGASK) (Fig. 1). This conserved AMP-binding site also appears in long-chain fatty acid-CoA ligases and 4-coumarate-CoA ligases, according to the InterPro web site. OsACOS12 also carries the conserved AMP-binding site (197-LPYSSGTTGVSK-208), whereas ACOS5 carries a similar amino acid sequence (186-LPFSSGTTGLQK-197), which was not determined to be IPR020845. As the result of these genetic analyses, *CjACOS5a* and *CjACOS5b* were thought to be sugi orthologs of *ACOS5* and *OsACOS12* and encode an acyl-CoA synthetase. The expression of the *CjACOS5a* gene in male strobili was about 280-fold higher than that detected in stems, whereas that in the leaves and female strobili was not significantly different from that in stems (Supplementary Fig. S1). Similarly, *CjACOS5b* was expressed most highly in male strobili, and their mRNA level was about 150-fold higher than that in stems. Accordingly, we expected that the *CjACOS5* genes mainly worked in sugi male strobili and was involved in pollen formation, similar to *Arabidopsis ACOS5* and rice *OsACOS12*.

**Figure 1.**
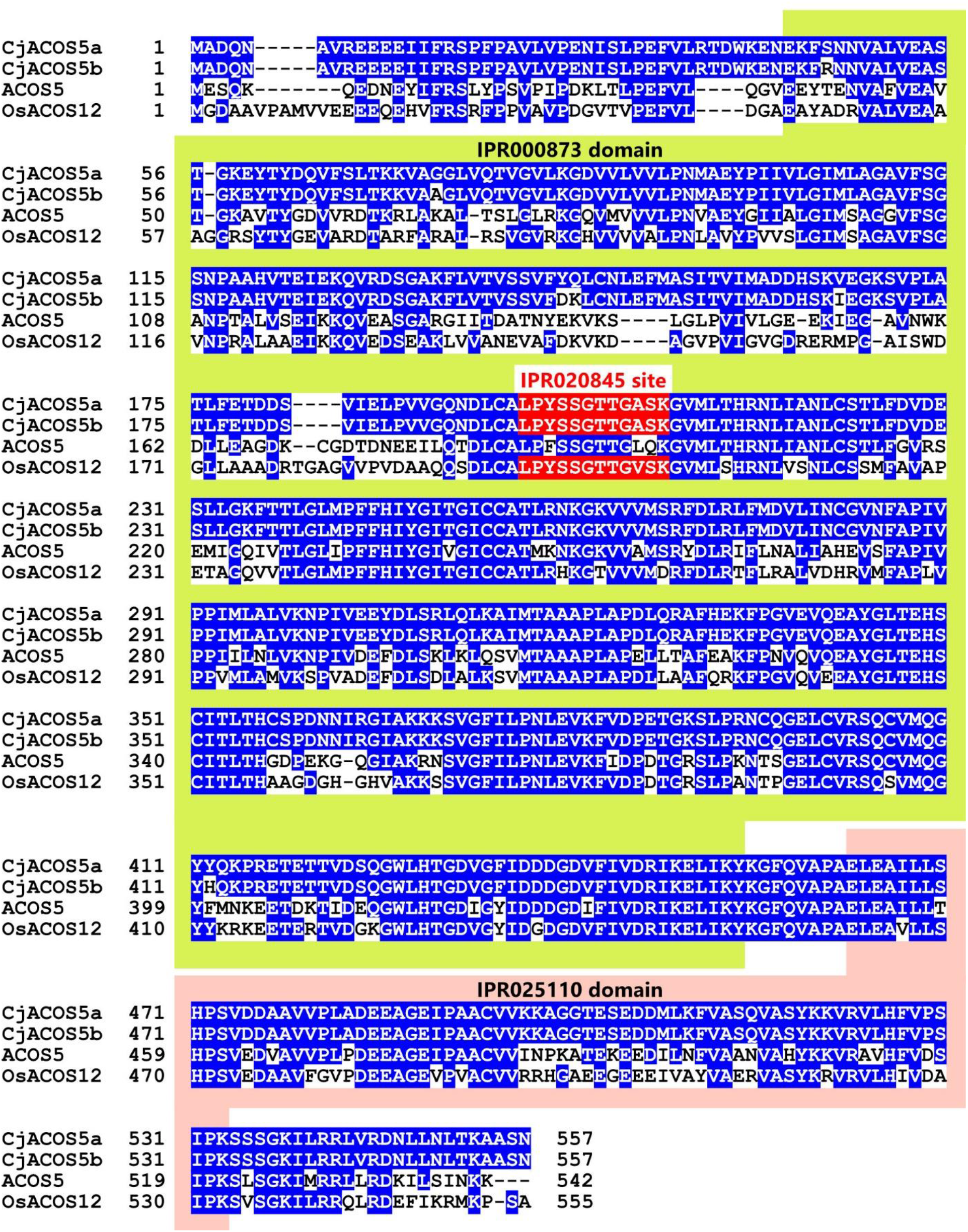
Comparison of the amino acid sequences of ACOS5 orthologs. CjACOS5a and CjACOS5b from sugi, ACOS5 from *Arabidopsis thaliana*, and OsACOS12 from rice were aligned using by the MAFFT program^75^. Identical amino acids to CjACOS5a are indicated in blue. The AMP-dependent synthetase/ligase domain (IPR000873), C-terminal domain in AMP-binding enzymes (IPR025110), and conserved AMP-binding site (IPR020845) are shown in green, pink, and red colors, respectively.

### CRISPR/Cas9 vector construction and sugi transformation

To induce loss of function of *CjACOS5* in sugi, we planned the target sites for DNA breaks by the CRISPR/Cas9 system. The recruitment of the complex of guide RNA and *Streptococcus pyogenes* Cas9 requires 5’-NGG-3’ as a protospacer adjacent motif (PAM) and the upstream target sequence. The AGG sequence located downstream of the start codon of *CjACOS5* was choose as a PAM (Fig. 2a). If the *CjACOS5* gene is broken at the predicted site, at 3 base pairs upstream of the PAM, short CjACOS5 proteins with about 34 or 24 amino acid residues would probably be synthesized by a flame shift. Two allelic *CjACOS5* genes, *CjACOS5a* and *CjACOS5b*, were distinguished by a single nucleotide polymorphism (T or G) located upstream of the start codon (Fig. 2a).

**Figure 2.**
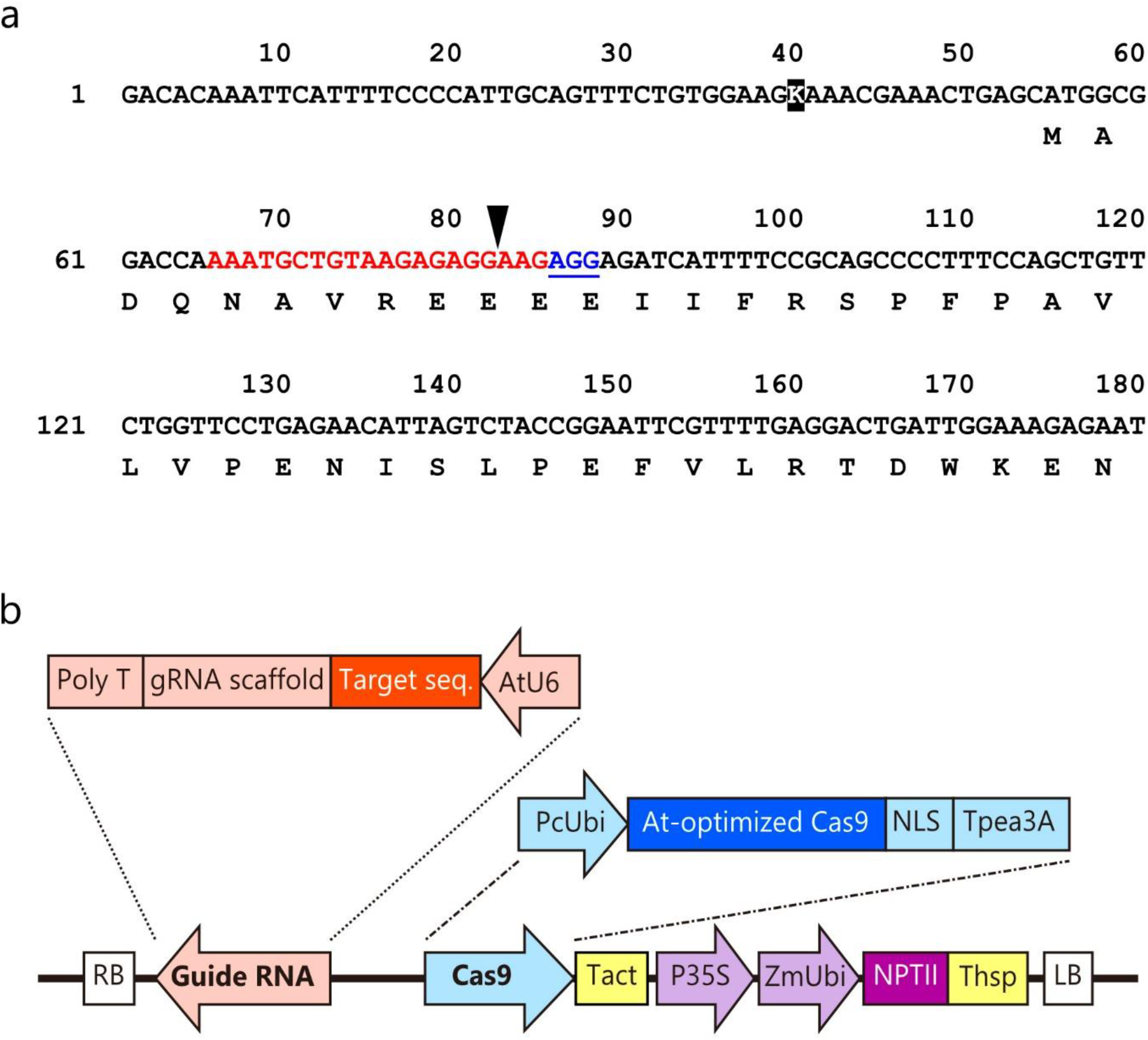
Design of the binary vector for the CRISPR/Cas9 system. (a) *CjACOS5* cDNA sequence neighboring the DNA break site for the CRISPR/Cas9 system. The target sequence and PAM are shown in red and underlined blue, respectively. The arrow head is located at a predicted DNA break site. The reverse-typed K is a single nucleotide polymorphism and indicates thymine in the *CjACOS5a* gene or guanine in the *CjACOS5b* gene. Amino acid residues are indicated under the cDNA sequence. (b) Schematic representation of the constructed CRISPR/Cas9 binary vector. This vector was named pBFGE1 and was derived from pZK_gYSA_FFCas9^37^. Twenty base pairs upstream of the PAM of *CjACOS5* were inserted as the target sequence. AtU6, *Arabidopsis thaliana* U6 promoter; gRNA scaffold, guide RNA; PcUbi, *Petroselinum crispum* ubiquitin promoter; At-optimized Cas9, *A. thaliana-* optimized *Streptococcus pyogenes* Cas9; NLS, simian virus 40 nuclear localization signal; Tpea3A, *Pisum sativum rbcS3A* terminator; Tact, *Oryza sativa* actin terminator; P35S, cauliflower mosaic virus 35S promoter; ZmUbi, *Zea mays* ubiquitin promoter; NPTII, neomycin phosphotransferase II; Thsp, *O. sativa* heat shock protein terminator; RB, right border; LB, left border.

A CRISPR/Cas9 vector was constructed for transformation of sugi. The original pZK_gYSA_FFCas9 vector was generated for dicots and is composed of the *A. thaliana* U6 promoter-driven guide RNA, the parsley ubiquitin promoter-driven Cas9, and the cauliflower mosaic virus (CaMV) 35S promoter-driven *NPTII* gene^37^. We modified pZK_gYSA_FFCas9 to generate the pBFGE1 vector, in which the *Zea mays* ubiquitin promoter was inserted between the CaMV 35S promoter and the *NPTII* gene, to further express the *NPTII* gene (Fig. 2b). The ubiquitin promoter has been reported to work well in sugi^38^. The 20-bp target sequence of *CjACOS5* was integrated into the guide RNA gene of the pBFGE1 vector, to break the *CjACOS5* genes (Fig. 2b).

The sugi embryogenic cell line #13-8-2 was infected with *Agrobacterium tumefaciens* GV3101 harboring the constructed pBFGE1 vector to generate transgenic sugi trees. After infection, the cells were cultivated in selection medium containing kanamycin and formed calli harboring the vector. Among the proliferated calli, three kanamycin-resistant calli lines were selected using PCR of the *NPTII* gene, and were designated as GE#1, GE#2 and GE#3 (Supplementary Fig. S2a). These kanamycin-resistant cells and the non-transgenic #13-8-2 cells were transferred to kanamycin-free maturation medium and somatic embryos were generated. Subsequently, the somatic embryos were moved onto germination media, germinated, and grown. The rooted transgenic sugi plantlets were planted in a pot and grown in a closed glass greenhouse (phytotron). All sugi plantlets regenerated from the GE#1, GE#2 and GE#3 calli possessed the *NPTII* gene in their leaves, whereas non-transgenic sugi plantlets obtained from the #13-8-2 cells did not (Supplementary Fig. S2b).

### Induced mutation of *CjACOS5*

To confirm the mutation of *CjACOS5* in the transgenic trees regenerated from the GE#1, GE#2 and GE#3 calli, genomic DNA was isolated from the leaves of each transgenic sugi plantlet, then DNA fragments including the target site of *CjACOS5* were amplified using PCR and cloned into a plasmid vector in *E. coli*. The plasmids were individually sequenced and analyzed. As a result, several deletion or insertion mutants were detected in the target DNA sequence of *CjACOS5a* and *CjACOS5b* in the transgenic sugi trees, whereas non-transgenic sugi trees that were regenerated from embryogenic cells had no mutation in the two genes (Supplementary Table S2).

The two GE#1 callus-derived transgenic sugi lines, GE#1-17 and GE#1-18, were chimeric plants (Fig. 3). The GE#1-17 sugi tree carried a 3-bp deletion and a 1-bp deletion of *CjACOS5a* and two types of 1-bp deletion of *CjACOS5b*. GE#1-18 contained wild-type and 1 bp-deleted *CjACOS5a* and a 1-bp deletion of *CjACOS5b*. In contrast, all eight transgenic sugi lines derived from GE#2 calli (GE#2-1 to GE#2-22) had the same biallelic mutation with a 1-bp deletion at the same position of *CjACOS5a* and *CjACOS5b* (Fig. 3 and Supplementary Table S2). These deletions of a single base pair (G:C) suggested the production of a truncated and mutated CjACOS5 protein of 34 amino acid residues (<u>MADQNAVRE</u>KRRSFSAAPFQLFWFLRTLVYRNSF). In this mutant CjACOS5 protein, the nine underlined-N-terminal amino acid residues alone are in accordance with the original CjACOS5 protein. Conversely, GE#3-derived transgenic trees include several types of mutation. Five transgenic sugi lines (GE#3-7, GE#3-8, GE#3-9, GE#3-11, and GE#3-12) were chimeric plants including wild-type and deleted *CjACOS5* genes (Fig. 3 and Supplementary Table S2). There were not only 1-bp deletions in those plants, but also a 45-bp deletion in GE#3-9, and a new DNA sequence comprising a 17-bp deletion and a 24-bp insertion in GE#3-12. GE#3-10 alone did not contain the wild-type *CjACOS5a* and *CjACOS5b* genes; thus, it was a biallelic mutant, similar to the GE#2 lines (Fig. 3). These results confirmed that the CRISPR/Cas9 system worked in sugi cells and induced mutations of the *CjACOS5a* and *CjACOS5b* genes.

**Figure 3.**
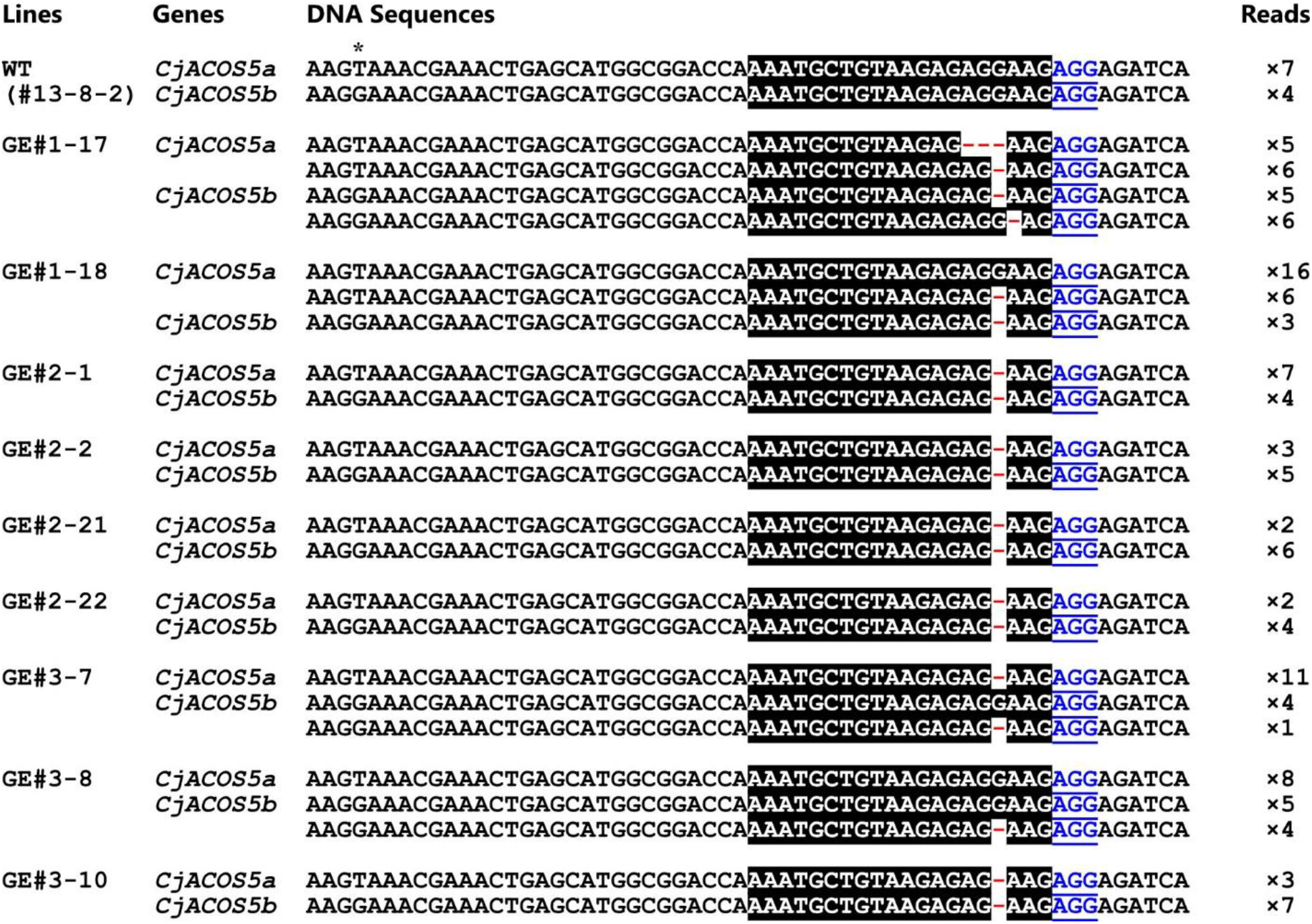
Mutations of *CjACOS5* genes by CRISPR/Cas9. DNA sequence of PCR-amplified DNAs neighboring the break site of *CjACOS5* in a non-transgenic sugi tree (WT) and in transgenic tree (GE#1, GE#2, and GE#3) lines. The asterisk shows a single nucleotide polymorphism in *CjACOS5a* and *CjACOS5b*. The target sequence and PAM are shown in reversed black and underlined blue, respectively. The gaps (-) represent deleted nucleotides. Reads mean the number of sequenced DNAs after cloning of the PCR products amplified using the leaf DNA.

### Morphogenesis and pollen production in *CjACOS5*-mutataed sugi trees

The transgenic sugi trees were planted in a pot and grown under the same conditions of outside temperature and sunshine in the phytotron (Supplementary Fig. S3a). The shape of the transgenic sugi trees from GE#2 lines and GE#3 lines appeared to be the same as that of non-transgenic sugi trees (Supplementary Fig. S3b, e–i). The initial height of trees from GE#2 lines and GE#3 lines was similar to that of non-transgenic sugi trees at 6 months, whereas GE#1-17 and GE#1-18 trees were significantly shorter than non-transgenic trees (Supplementary Fig. S3c, d, j). Unfortunately, GE#1-17 and GE#1-18 trees withered and died in the winter of 2017, probably because of severe growth suppression.

Flowering of sugi can be induced by spraying a gibberellin A3 (GA_3_) solution onto the twigs with leaves^39^. At the end of July/beginning of August of 2017, the young transgenic sugi trees with the mutation of *CjACOS5* and non-transgenic sugi trees were treated twice with GA_3_ and cultivated in the phytotron. Visible flower buds appeared in October and developed to male strobili. In January of 2018, yellow-powdered pollen was packed in the male strobili of non-transgenic sugi trees (Fig. 4a). Spherical pollen grains in the male strobili were detected under a microscope.

**Figure 4.**
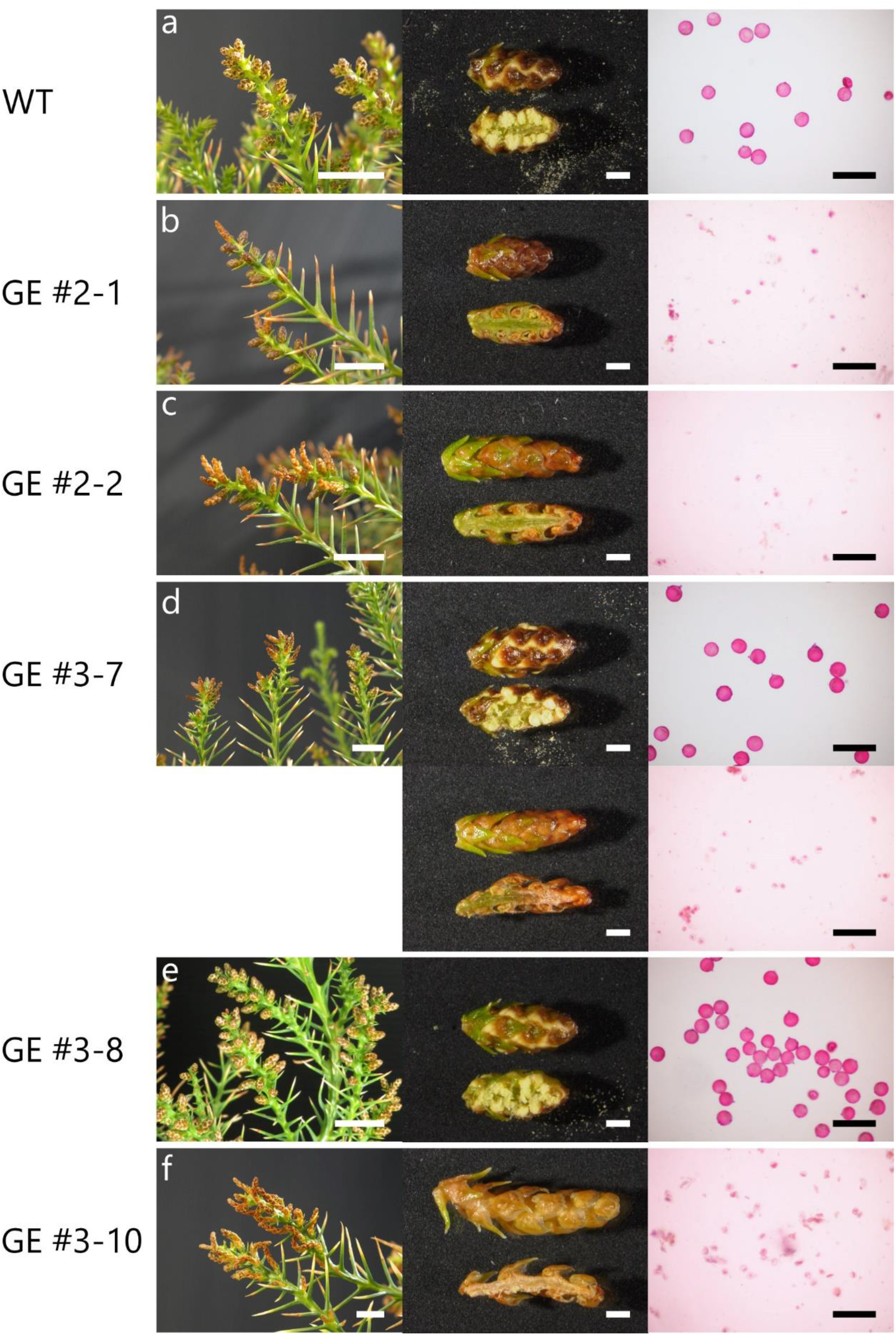
Male strobili of genome-edited sugi trees in 2018. (a) Non-transgenic sugi (WT) male strobili (left; scale bar, 1 cm), magnified male strobili (middle; scale bar, 1 mm), and razor-cutting suspension of male strobili (right; scale bar, 100 μm). The magnified strobili are shown as an outside (top) and a vertical section (bottom). Spherical pollen grains and the suspension were stained with Calberla’s fuchsin staining solution (right). (b) Male strobili of the genome-edited GE#2-1 sugi tree, a biallelic mutant of *CjACOS5*. (c) GE#2-2, a biallelic mutant. (d) GE#3-7, a chimeric transgenic sugi tree. Normal male strobili and no-pollen male strobili are indicated. (e) GE#3-8, a chimeric transgenic sugi tree with pollen. (f) GE#3-10, a biallelic mutant without pollen. The photographs were taken in January, 2018.

In contrast, GE#2 sugi trees from GE#2-1 to GE#2-6, which were biallelic mutants of *CjACOS5a* and *CjACOS5b*, produced male strobili without pollen grains (Fig. 4b, c, and Supplementary Table S2). The male strobili were finely sliced using a razor blade in water, and the suspension was stained and observed under a microscope, however, no-pollen grains were detected and small cells were observed. Conversely, GE#3 sugi trees resulted in three types, as follows. The chimeric trees with wild-type and deleted genes, GE#3-7, GE#3-9, GE#3-11, and GE#3-12, had pollen-including male strobili and no-pollen male strobili (Fig. 4d and Supplementary Table S2). GE#3-8 was also a chimeric tree but had male strobili with pollen (Fig. 4e). GE#3-10, the sole biallelic mutant among the GE#3 lines, had no-pollen at all, similar to the GE#2 lines (Fig. 4f). Therefore, disruption of both *CjACOS5a* and *CjACOS5b* led to the loss of the ability of pollen production in male strobili. Moreover, the pollen productivity was shown to be maintained exclusively by one allele gene, *CjACOS5a* or *CjACOS5b*.

A sterile male strobilus of GE#2-1 and GE#2-2 tree carried only deleted *CjACOS5* genes as same as their leaves (Supplementary Fig. S4). Conversely, a sterile male strobilus of GE#3-7 carried not only deleted *CjACOS5* genes but also wild-type genes similar to a fertile male strobilus of GE#3-7. That is probably because a male strobilus had been constructed with not only mutated reproductive cells but also somatic cells without mutation. Additionally, the male strobilus of GE#3-7 possessed different types of mutations, a 2-bp deletion and a 4-bp deletion, which were not been observed in the leaves (Supplementary Fig. S4 and Fig. 3). The plausible reason is that GE#3-7 is a chimeric tree carrying the active *Cas9* gene which has been inducing the mutation of the *CjACOS5* genes.

To determine the reproducibility of the no-pollen flowers, the mutated sugi trees were again treated with GA_3_ in the summer of 2018. They were grown in the phytotron and in a special netted-house. Biallelic mutant sugi trees, such as GE#2-1 and GE#3-10, grown in the phytotron had male strobili but did not produce pollen in 2019 (Supplementary Fig. S5b, c, f). The chimeric mutants (GE#3-7, GE#3-9, GE#3-11, and GE#3-12) produced two types of strobili, i.e., with or without pollen (Supplementary Fig. S5d). GE#3-8 produced only fertile male strobili similar to the result of the preceding year (Supplementary Fig. S5e). The special netted-house is bigger than the phytotron and has a glass roof and a metal mesh for ventilation.

That did not have an air conditioner, so that its temperature depended on the outside temperature and sunlight but often reached around 40°C in the summer. The sugi trees suffered no damage under high temperature conditions. In the special netted-house, the biallelic mutant trees (GE#2-21 and GE#2-22) did not produce pollen in their male strobili, whereas the non-transgenic trees had pollen (Supplementary Fig. S5g–i). During the third experimental period, from 2019 to 2020, *CjACOS5* biallelic mutant GE#2 lines also had male strobili but did not produce pollen in the special netted-house (Supplementary Fig. S6). Accordingly, it was suggested that CRISPR/Cas9-induced mutation of *CjACOS5* genes stably maintained the no-pollen phenotype over a few years in sugi trees.

## Discussion

Pollen-free sugi trees have been expected to mitigate Japanese cedar pollinosis. For that reason, male-sterile pollen-free sugi trees have been the goal of time-consuming and laborious research, and are rarely found in the field or in the test forests. In the present study, we accomplished the artificial generation of pollen-free sugi using genome editing technology. Sugi *CjACOS5* genes were targeted and mutated by a CRISPR/Cas9 vector via *Agrobacterium-* mediated transformation. The regenerated sugi trees with biallelic *CjACOS5* deletions could not produce pollen in their male strobili (Fig. 4). Male sterility with no-pollen has been regarded as being important and useful for human health, the economy, biodiversity, and agriculture. For example, if plant species with a large amount of allergenic pollen, such as sugi, cypress, and birch trees and pasture grasses, cannot produce pollen, the number of pollen-allergic patients will decrease. As the result, the pollinosis-related health expenditures and economic loss will decrease^40^. Pollen-free plants can also avoid unnecessary gene flow, which causes problems of biodiversity in the case of invasive and GMO plants^41^. Male sterility in a variety of crops (maize, rice, wheat, etc.) has helped produce hybrid seeds more efficiently than before^42–44^. Using genome editing technology, it is possible to turn a plant species with pollen into pollen-free varieties, even if wild pollen-free mutants had never been found in the same plant species. Thus, the generation of male-sterile plants using the CRISPR/Cas9 system is innovative and greatly available.

The no-pollen phenotype in male strobili of the genome-edited sugi trees was thought to be caused by the mutation of *CjACOS5a* and *CjACOS5b*, as demonstrated by the results obtained for different mutation types. The GE#2 lines and the GE#3-10 line had biallelic mutation of both deleted *CjACOS5a* and *CjACOS5b*, and they showed a no-pollen phenotype in male strobili (Fig. 3 and Fig. 4). These results demonstrated that biallelic mutation of *CjACOS5a* and *CjACOS5b* was closely related to the no-pollen phenotype. GE#3-7 produced male strobili partly with pollen or without pollen. Because GE#3-7 was a chimeric tree carrying the deleted *CjACOS5a* gene and the normal and delated *CjACOS5b* gene, it was suggested that normal *CjACOS5b* functioned in pollen production. Male strobili with pollen and without pollen were thought to be derived from biallelic mutant cells and monoallelic mutant cells, respectively. Conversely, it was thought that *CjACOS5a* alone could lead to the production of pollen, because GE#3-8 had normal *CjACOS5a*, deleted *CjACOS5b*, and normal *CjACOS5b* and produced only male strobili with pollen. Therefore, it was concluded that the no-pollen phenotype observed in this study was caused by loss of function of both the *CjACOS5a* and *CjACOS5b* genes, and that one allelic gene, alone, either *CjACOS5a* or *CjACOS5b*, was essential to generate pollen in male strobili. If possible, to confirm that the no-pollen phenotype was not caused by off-target mutagenesis, we should elucidate the existence of off-target mutagenesis in the genome-edited sugi trees in the same way as that used for several model plants^45^. However, the off-target mutagenesis in sugi is difficult to investigate because its genomic DNA sequence has never been adequately read. For the development of the genome editing technology of sugi, investigation of the off-target mutation is surely needed in the future.

The sugi *CjACOS5* gene was revealed to be essential for pollen production in this study. The *Arabidopsis* ACOS5 protein is an acyl-CoA synthetase that is estimated to be linked to the biosynthesis of sporopollenin in the exine of pollen^12^. The rice OsACOS12 protein is the ortholog of ACOS5 and catalyzes the condensation of oleic acid and CoA *in vitro*^14^. Although the physiological and biochemical functions of CjACOS5 were not clarified in this study, CjACOS5 was deduced to function as an acyl-CoA synthetase based on the structural analogy to ACOS5 and OsACOS12 (Fig. 1). Sporopollenin biosynthesis requires not only acyl-CoA synthetase, but also several other enzymes encoded by the *MALE STERILITY 2* (*MS2*)^46,47^, *CYP703A2^48^, CYP704B1^49^, LESS ADHESIVE POLLEN 3 (LAP3)^50^, LAP5/POLYKETIDE SYNTHASE B* (*PKSB*)^51,52^, *LAP6*/*PKSA*^51,52^, and *DIHYDROFLAVONOL 4-REDUCTASE-LIKE 1 (DRL1)/TETRAKETIDE α-PYRONE REDUCTASE 1 (TKPR1*) genes^53,54^. The *Arabidopsis ms2* mutant, the *drl1/tkpr1* mutant, the *lap3* mutant, and the double mutant of *lap5* and *lap6 (pksb* and *pksa*) show complete male sterility. Similarly, the rice *defective pollen wall (dpw*, the ortholog of *MS2*)^55^, *Ospks1*^56^, *Ospks2*^57^, and *Ostkpr1*^58^ mutants exhibit a male-sterile phenotype individually. The wheat *TaNP1* gene (encoding a putative glucose-methanol-choline oxidoreductase) is an ortholog of *OsNP1* that is involved in tapetum degeneration and pollen exine formation^59^. As the result of *TaNP1* disruption by CRISPR/Cas9, triple homozygous mutants of *TaNP1* showed complete male sterility. The tomato *SlSTR1* gene (a stamen-specific expressed strictosidine synthase) is an ortholog of *Arabidopsis LAP3*, and CRISPR/Cas9-mediated mutagenesis of *SlSTR1* confers no viability on pollen^60^. Those male sterility-related gene orthologs of sugi probably can be utilized in addition to *CjACOS5* to generate new varieties with male sterility by CRISPR/Cas9.

Our study established that the genome editing technology is available as a method for generating pollen-free sugi trees. So far, conventional crossbreeding programs using wild male-sterile sugi mutants have been useful for the generation of new pollen-free varieties of sugi. Because these crossed varieties can be freely handled, unlike the case of GMO trees, the new pollen-free sugi varieties that are the progeny of sugi plus trees and wild pollen-free mutants have been gradually planted in Japan^3^. In contrast, the *CjACOS5*-disrupted pollen-free sugi trees obtained in this study are GMO trees and, thus, are currently restricted for handling and planting by law. However, the genome editing technology, including CRISPR/Cas9, has several advantages compared with conventional tree breeding, as follows. First, if we could identify target genes involved in excellent traits, genome editing technology would be able to directly modify the target genes and generate their mutant trees in cases in which mutants of the target genes are not found in nature. Second, CRISPR/Cas9 can edit several target genes simultaneously and, thus, can modify multiple traits. Considering the combination of expected traits, such as no-pollen, fast growth, quality of wood, straightness, and stress tolerance, multiple values can be conferred to sugi breeding using multiplex gene editing technology. Last, the genome editing technology can not affect every gene other than the target genes, if off-target mutagenesis can be controlled fully. Therefore, the genome editing technology is difficult to perturb the genetic composition of well-established varieties unlike crossbreeding. Recently, several methods have been developed for the generation of non-transgenic genome-edited plants^61–63^. Consequently, it is expected that they will be utilized in combination with the advantage of genome editing technology and conventional breeding methods for tree breeding.

## Methods

### Vector construction

pZK_gYSA_FFCas9 vector^37^ was digested with XbaI, the recognition site of which was located between the cauliflower mosaic virus 35S promoter and the *NPTII* gene, and blunted with Klenow fragment. About 2 kb of *Zea mays* ubiquitin promoter was inserted into the blunted pZK_gYSA_FFCas9 vector^64^. The resulting binary vector was designated as pBFGE1. Two synthetic DNAs (Supplementary Table S1) were annealed and ligated into the BbsI-digested pUC19_AtU6oligo vector^65^. A derivative vector was digested with I-SceI, and DNA fragments including the oligonucleotides were excised. The DNA fragments were inserted into I-SceI digested pBFGE1.

### Plant materials and sugi transformation

The sugi (*Cryptomeria japonica* D. Don) embryonic cell line #13-8-2 was induced from an immature seed using previously described methods^66^. The #13-8-2 cells were transformed using *A. tumefaciens* GV3101 harboring the constructed pBFGE1 vector as described previously^67,68^. Selected kanamycin-resistant calli were regenerated through somatic embryogenesis^69^. Regenerated sugi plantlets were acclimatized in moist peat moss and Kanuma pumice, then transferred to moist vermiculite in a pot. They were cultivated under natural-day-length conditions in a closed glass greenhouse (phytotron) or a special netted-house. The temperature inside the phytotron was controlled by an air conditioner following the outdoor temperature. The special netted-house was a greenhouse with metal mesh windows that passed the outside air but impeded invasion by insects.

### DNA analysis

Orthologs of the CMFL003_A04 cDNA were searched using the blastn program in the latest NCBI nucleotide database^70,71^. Total RNA was prepared as described previously^72^. The cDNA cloning and expression analysis of *CjACOS5* genes are performed as described in Supplementary Information. A functional domain analysis of ACOS5-related protein sequences was performed on the InterPro website^73^. The *NPTII* and *CjACOS5* genes in the transgenic sugi trees were confirmed using PCR (Supplementary Information). To elucidate the mutation induced by genome editing, a DNA region including the predicted DNA break site for the CRISPR/Cas9 system was amplified using PCR and sequenced (Supplementary Information).

### Induction and observation of male strobili

To induce flowering, 100 mg/L of a gibberellin A3 (GA_3_) aqueous solution was sprayed on sugi plantlets. In the first experiment, GA_3_ treatments were performed twice, on July 24 and August 3, 2017. Male strobili were observed in January, 2018. Pollen and the suspension of sliced male strobili were stained with Calberla’s fuchsin staining solution, for microscopy^74^. In the second and third experiments, GA_3_ treatments were performed in late July, 2018 and 2019, respectively. Thereafter, male strobili and pollen were analyzed in the following year in the same way as that described for first experiment.

## Acknowledgments

The authors are grateful to Ms. K. Nemoto, Ms. S. Tanaka, and Ms. A. Hagiwara for helpful assistance. This work was partly supported by Council for Science, Technology and Innovation (CSTI), Cross-ministerial Strategic Innovation Promotion Program (SIP), “Technologies for creating next-generation agriculture, forestry and fisheries” (funding agency: Bio-oriented Technology Research Advancement Institution, NARO), and by JSPS KAKENHI Grant Number JP20H03037.

## Author contributions

NF and MN designed the study and constructed the vectors. ME, MM, ST and SIK were concerned in the construction of the vectors. YO, KK, YN, and TT were involved in the transformation of sugi. TEM regenerated the transgenic trees and treated them with gibberellin. MN analyzed the transgenes, flowers and pollen, and wrote the manuscript. All authors provided critical feedback for the manuscript.

## Data availability

The data that support the findings of this study are available from the corresponding author upon reasonable request.

## Additional information

### Supplementary Information

The online version contains supplementary material.

## References

1 Christenhusz, M. J. M. et al. A new classification and linear sequence of extant gymnosperms. Phytotaxa 19, 55–70 (2011).

2 Forestry Agency. Statistical Handbook of Forest and Forestry. (Japan Forest Foundation, 2017) (in Japanese).

3 Forestry Agency. Annual Report on Forest and Forestry in Japan. Fiscal Year 2018 (Summary). (Ministry of Agriculture, Forestry, and Fisheries, 2019).

4 Fujimura, T. & Kawamoto, S. Spectrum of allergens for Japanese cedar pollinosis and impact of component-resolved diagnosis on allergen-specific immunotherapy. Allergol. Int. 64, 312–320 (2015).

5 Horiguchi, S. & Saito, Y. Japanese cedar pollinosis in Nikko, Japan. Arerugi 13, 16–18 (1964) (in Japanese with English abstract).

6 Nakamura, A., Asai, T., Yoshida, K., Baba, K. & Nakae, K. Allergic rhinitis epidemiology in Japan. J. Otolaryngol. Jpn. 105, 215–224 (2002) (in Japanese with English abstract).

7 Okuda, M. Epidemiology of Japanese cedar pollinosis throughout Japan. Ann. Allergy Asthma Immunol. 91, 288–296 (2003).

8 Baba, K. & Nakae, K. Epidemiology of nasal allergy through Japan in 2008. Prog. Med. 28, 2001–2012 (2008) (in Japanese).

9 Matsubara, A. et al. Epidemiological survey of allergic rhinitis in Japan 2019. J. Otolaryngol. Jpn. 123, 485–490 (2020) (in Japanese with English abstract).

10 Vedel, F. et al. Molecular basis of nuclear and cytoplasmic male sterility in higher plants. Plant Physiol. Biochem. 32, 601–618 (1994).

11 Horner, H. & G. Palmer, R. Mechanisms of genic male sterility. Crop Sci. 35, 1527–1535 (1995).

12 de Azevedo Souza, C. et al. A novel fatty acyl-CoA synthetase is required for pollen development and sporopollenin biosynthesis in *Arabidopsis*. Plant Cell 21, 507–525 (2009).

13 Li, Y. et al. OsACOS12, an orthologue of *Arabidopsis* acyl-CoA synthetase5, plays an important role in pollen exine formation and anther development in rice. BMC Plant Biol. 16, 256 (2016).

14 Yang, X. et al. Rice fatty acyl-CoA synthetase OsACOS12 is required for tapetum programmed cell death and male fertility. Planta 246, 105–122 (2017).

15 Taira, H., Teranishi, H. & Kenda, Y. A case study of male sterility in sugi (*Cryptomeria japonica*). J. Jpn. For. Soc. 75, 377–379 (1993) (in Japanese with English abstract).

16 Saito, M., Taira, H. & Furuta, Y. Cytological and genetical studies on male sterility in *Cryptomeria japonica* D. Don. J. For. Res. 3, 167–173 (1998).

17 Yoshii, E. & Taira, H. Cytological and genetical studies on male sterile sugi (*Cryptomeria japonica* D. Don), Shindai 1 and Shindai 5. J. Jpn. For. Soc. 89, 26–30 (2007) (in Japanese with English abstract).

18 Ueuma, H., Yoshii, E., Hosoo, Y. & Taira, H. Cytological study of a male-sterile *Cryptomeria japonica* that does not release microspores from tetrads. J. For. Res. 14, 123–126 (2009).

19 Miyajima, D., Yoshii, E., Hosoo, Y. & Taira, H. Cytological and genetic studies on male sterility in *Cryptomeria japonica* D. Don (Shindai 8). J. Jpn. For. Soc. 92, 106–109 (2010) (in Japanese with English abstract).

20 Moriguchi, Y. et al. The construction of a high-density linkage map for identifying SNP markers that are tightly linked to a nuclear-recessive major gene for male sterility in *Cryptomeria japonica* D. Don. BMC Genomics 13, 95 (2012).

21 Moriguchi, Y. et al. Establishment of a microsatellite panel covering the sugi (*Cryptomeria japonica*) genome, and its application for localization of a male-sterile gene (*ms-2*). Mol. Breed. 33, 315–325 (2014).

22 Moriguchi, Y. et al. A high-density linkage map with 2560 markers and its application for the localization of the male-sterile genes *ms3* and *ms4* in *Cryptomeria japonica* D. Don. Tree Genet. Genomes 12, 57 (2016).

23 Mishima, K. et al. Identification of novel putative causative genes and genetic marker for male sterility in Japanese cedar (*Cryptomeria japonica* D.Don). BMC Genomics 19, 277 (2018).

24 Hasegawa, Y. et al. Fine mapping of the male-sterile genes (*MS1, MS2, MS3*, and *MS4*)and development of SNP markers for marker-assisted selection in Japanese cedar (*Cryptomeria japonica* D. Don). PLoS One 13, e0206695 (2018).

25 Hasegawa, Y. et al. Identification and genetic diversity analysis of a male-sterile gene (*MS1*) in Japanese cedar (*Cryptomeria japonica* D. Don). Sci. Rep. 11, 1496 (2021).

26 Saito, M., Koga, Y., Furuta, Y. & Taira, H. Selections of male sterile sugi (*Cryptomeria japonica* D. Don) trees from open pollinated seedlings in a seed orchard. J. Jpn. For. Soc. 87, 1–7 (2005) (in Japanese with English abstract).

27 Jia, H. & Wang, N. Targeted genome editing of sweet orange using Cas9/sgRNA. PLoS One 9, e93806 (2014).

28 Nishitani, C. et al. Efficient genome editing in apple using a CRISPR/Cas9 system. Sci. Rep. 6, 31481 (2016).

29 Ren, C. et al. CRISPR/Cas9-mediated efficient targeted mutagenesis in Chardonnay (*Vitis vinifera* L.). Sci. Rep. 6, 32289 (2016).

30 Fan, D. et al. Efficient CRISPR/Cas9-mediated targeted mutagenesis in *Populus* in the first generation. Sci. Rep. 5, 12217 (2015).

31 Zhou, X., Jacobs, T. B., Xue, L. J., Harding, S. A. & Tsai, C. J. Exploiting SNPs for biallelic CRISPR mutations in the outcrossing woody perennial *Populus* reveals 4-coumarate:CoA ligase specificity and redundancy. New Phytol. 208, 298–301 (2015).

32 Nanasato, Y. et al. CRISPR/Cas9-mediated targeted mutagenesis in Japanese cedar (*Cryptomeria japonica* D. Don). Sci. Rep. 11, 16186 (2021).

33 Poovaiah, C. et al. Genome editing with CRISPR/Cas9 in *Pinus radiata* (D. Don). BMC Plant Biol. 21, 363 (2021).

34 Cui, Y. et al. Efficient multi-sites genome editing and plant regeneration *via* somatic embryogenesis in *Picea glauca*. Front. Plant Sci. 12, 751891 (2021).

35 Nagamine, A. & Ezura, H. Genome editing for improving crop nutrition. Front. Genome Ed. 4, 850104 (2022).

36 Futamura, N. et al. Characterization of expressed sequence tags from a full-length enriched cDNA library of *Cryptomeria japonica* male strobili. BMC Genomics 9, 383 (2008).

37 Mikami, M., Toki, S. & Endo, M. Comparison of CRISPR/Cas9 expression constructs for efficient targeted mutagenesis in rice. Plant Mol. Biol. 88, 561–572 (2015).

38 Taniguchi, T., Ohmiya, Y., Kurita, M., Tsubomura, M. & Kondo, T. Regeneration of transgenic *Cryptomeria japonica* D. Don after *Agrobacterium tumefaciens*-mediated transformation of embryogenic tissue. Plant Cell Rep. 27, 1461–1466 (2008).

39 Nagao, A., Sasaki, S. & Pharis, R. P. Cryptomeria japonica in CRC Handbook of Flowering, Volume IV (ed. Halevy, A. H.) 247–269(CRC Press, 1989).

40 Blaiss, M. S. Allergic rhinitis: direct and indirect costs. Allergy. Asthma. Proc. 31, 375–380 (2010).

41 Fritsche, S., Klocko, A. L., Boron, A., Brunner, A. M. & Thorlby, G. Strategies for engineering reproductive sterility in plantation forests. Front. Plant Sci. 9, 1671 (2018).

42 Wan, X. et al. Maize genic male-sterility genes and their applications in hybrid breeding: progress and perspectives. Mol. Plant 12, 321–342 (2019).

43 Abbas, A. et al. Exploiting genic male sterility in rice: from molecular dissection to breeding applications. Front. Plant Sci. 12, 629314 (2021).

44 Singh, M., Albertsen, M. C. & Cigan, A. M. Male fertility genes in bread wheat (*Triticum aestivum* L.) and their utilization for hybrid seed production. Int. J. Mol. Sci. 22, 8157 (2021).

45 Graham, N. et al. Plant genome editing and the relevance of off-target changes. Plant Physiol. 183, 1453–1471 (2020).

46 Aarts, M. G. et al. The *Arabidopsis MALE STERILITY 2* protein shares similarity with reductases in elongation/condensation complexes. Plant J. 12, 615–623 (1997).

47 Chen, W. et al. Male Sterile2 encodes a plastid-localized fatty acyl carrier protein reductase required for pollen exine development in Arabidopsis. Plant Physiol. 157, 842–853 (2011).

48 Morant, M. et al. CYP703 is an ancient cytochrome P450 in land plants catalyzing in-chain hydroxylation of lauric acid to provide building blocks for sporopollenin synthesis in pollen. Plant Cell 19, 1473–1487 (2007).

49 Dobritsa, A. A. et al. CYP704B1 is a long-chain fatty acid ω-hydroxylase essential for sporopollenin synthesis in pollen of *Arabidopsis*. Plant Physiol. 151, 574–589 (2009).

50 Dobritsa, A. A. et al. LAP3, a novel plant protein required for pollen development, is essential for proper exine formation. Sex. Plant Reprod. 22, 167–177 (2009).

51 Dobritsa, A. A. et al. LAP5 and *LAP6* encode anther-specific proteins with similarity to chalcone synthase essential for pollen exine development in Arabidopsis. Plant Physiol. 153, 937–955 (2010).

52 Kim, S. S. et al. LAP6/POLYKETIDE SYNTHASE A and *LAP5/POLYKETIDE SYNTHASE B* encode hydroxyalkyl α-pyrone synthases required for pollen development and sporopollenin biosynthesis in *Arabidopsis thaliana*. Plant Cell 22, 4045–4066 (2010).

53 Tang, L. K., Chu, H., Yip, W. K., Yeung, E. C. & Lo, C. An anther-specific dihydroflavonol 4-reductase-like gene (*DRL1*) is essential for male fertility in Arabidopsis. New Phytol. 181, 576–587 (2009).

54 Grienenberger, E. et al. Analysis of *TETRAKETIDE α-PYRONE REDUCTASE*function in *Arabidopsis thaliana* reveals a previously unknown, but conserved, biochemical pathway in sporopollenin monomer biosynthesis. Plant Cell 22, 4067–4083 (2010).

55 Shi, J. et al. DEFECTIVE POLLEN WALL is required for anther and microspore development in rice and encodes a fatty acyl carrier protein reductase. Plant Cell 23, 2225–2246 (2011).

56 Zou, T. et al. OsLAP6/OsPKS1, an orthologue of *Arabidopsis PKSA/LAP6*, is critical for proper pollen exine formation. Rice 10, 53 (2017).

57 Zhu, X. et al. The polyketide synthase OsPKS2 is essential for pollen exine and Ubisch body patterning in rice. J. Integr. Plant Biol. 59, 612–628 (2017).

58 Xu, D. et al. Ostkpr1 functions in anther cuticle development and pollen wall formation in rice. BMC Plant Biol. 19, 104 (2019).

59 Li, J., Wang, Z., He, G., Ma, L. & Deng, X. W. CRISPR/Cas9-mediated disruption of *TaNP1* genes results in complete male sterility in bread wheat. J. Genet. Genomics 47, 263–272 (2020).

60 Du, M. et al. A biotechnology-based male-sterility system for hybrid seed production in tomato. Plant J. 102, 1090–1100 (2020).

61 Metje-Sprink, J., Menz, J., Modrzejewski, D. & Sprink, T. DNA-free genome editing: past, present and future. Front. Plant Sci. 9, 1957 (2018).

62 Wada, N., Ueta, R., Osakabe, Y. & Osakabe, K. Precision genome editing in plants: state-of-the-art in CRISPR/Cas9-based genome engineering. BMC Plant Biol. 20, 234 (2020).

63 Imai, R. et al. In planta particle bombardment (iPB): a new method for plant transformation and genome editing. Plant Biotechnol. (Tokyo) 37, 171–176 (2020).

64 Christensen, A. H., Sharrock, R. A. & Quail, P. H. Maize polyubiquitin genes: structure, thermal perturbation of expression and transcript splicing, and promoter activity following transfer to protoplasts by electroporation. Plant Mol. Biol. 18, 675–689 (1992).

65 Ito, Y., Nishizawa-Yokoi, A., Endo, M., Mikami, M. & Toki, S. CRISPR/Cas9-mediated mutagenesis of the *RIN* locus that regulates tomato fruit ripening. Biochem. Biophys. Res. Commun. 467, 76–82 (2015).

66 Taniguchi, T., Konagaya, K. & Nanasato, Y. Somatic embryogenesis in artificially pollinated seed families of 2nd generation plus trees and cryopreservation of embryogenic tissue in *Cryptomeria japonica* D. Don (Sugi). Plant Biotechnol. (Tokyo) 37, 239–245 (2020).

67 Konagaya, K., Kurita, M. & Taniguchi, T. High-efficiency *Agrobacterium-mediated* transformation of *Cryptomeria japonica* D. Don by co-cultivation on filter paper wicks followed by meropenem treatment to eliminate *Agrobacterium*. Plant Biotechnol. (Tokyo) 30, 523–528 (2013).

68 Konagaya, K., Nanasato, Y. & Taniguchi, T. A protocol for *Agrobacterium-mediated* transformation of Japanese cedar, Sugi (*Cryptomeria japonica* D. Don) using embryogenic tissue explants. Plant Biotechnol. (Tokyo) 37, 147–156 (2020).

69 Maruyama, E. & Hosoi, Y. Polyethylene glycol enhance somatic embryo production in Japanese cedar (*Cryptomeria japonica* D. Don). Propag. Ornam. Plants 7, 57–61 (2007).

70 Zhang, Z., Schwartz, S., Wagner, L. & Miller, W. A greedy algorithm for aligning DNA sequences. J. Comput. Biol. 7, 203–214 (2000).

71 Sayers, E. W. et al. Database resources of the national center for biotechnology information. Nucleic Acids Res. 50, D20–D26 (2022).

72 Nishiguchi, M., Nanjo, T. & Yoshida, K. The effects of gamma irradiation on growth and expression of genes encoding DNA repair-related proteins in Lombardy poplar (*Populus nigra* var. italica). J. Environ. Radioact. 109, 19–28 (2012).

73 Jones, P. et al. InterProScan 5: genome-scale protein function classification. Bioinformatics 30, 1236–1240 (2014).

74 Ogden, E. C. & Raynor, G. S. A new sampler for airborne pollen: the rotoslide. J. Allergy 40, 1–11 (1967).

75 Nakamura, T., Yamada, K. D., Tomii, K. & Katoh, K. Parallelization of MAFFT for large-scale multiple sequence alignments. Bioinformatics 34, 2490–2492 (2018).

