## Supplementary Information for "CRISPR/Cas9-mediated disruption of *CjACOS5* confers no-pollen formation on sugi trees (*Cryptomeria japonica* D. Don)"

#### **Supplementary methods**

For the cDNA cloning of *CjACOS5a* and *CjACOS5b*, total RNA was isolated from male strobili of sugi #13-8-2 line as described previously[1]. First strand cDNA was synthesized using AffinityScript QPCR cDNA Synthesis Kit (Agilent Technologies). Using this first strand cDNA, the *CjACOS5* cDNAs were amplified by PCR with Q5 Hot Start High-Fidelity DNA Polymerase (New England Biolabs). The PCR primers used here are shown in Supplementary Table S1. The PCR products were cloned into the pBlueScript II vector and sequenced.

For gene expression analysis, the investigated organs were sampled from a non-transgenic sugi tree in December and frozen in liquid nitrogen. Total RNA preparation and reverse transcription quantitative real-time PCR (RT-qPCR) were performed as described previously[1]. The RT-qPCR primers are shown in Supplementary Table S1. The relative mRNA level was normalized to the mRNA level of the eukaryotic translation initiation factor 1 (*CjeIF*) gene[2].

The *NPTII* and *CjACOS5* genes in the transformed cells and the regenerated trees were confirmed using PCR and 0.7% agarose gel electrophoresis. Genomic DNA was isolated from embryogenic cells and from leaves using the DNeasy Plant Mini Kit (Qiagen). PCR was performed using Q5 Hot Start High-Fidelity DNA Polymerase and the PCR primers as shown in Supplementary Table S1. The amplified *CjACOS5* DNA fragments were cloned into the pBlueScript II vector and sequenced for analysis of mutation.

For statistical analysis, the Tukey–Kramer test and Dunnett's test were performed using the KaleidaGraph software Ver. 4.5 (Synergy Software).

### References

1 Nishiguchi, M., Nanjo, T. & Yoshida, K. The effects of gamma irradiation on growth and expression of genes encoding DNA repair-related proteins in Lombardy poplar (*Populus nigra* var. *italica*). *J. Environ. Radioact.* **109**, 19-28 (2012).

2 Katahata, S-I., Futamura, N., Igasaki, T. & Shinohara, K. Functional analysis of *SOC1*-like and *AGL6*-like MADS-box genes of the gymnosperm *Cryptomeria japonica*. *Tree Genet. Genomes* **10**, 317-327 (2014).

### Figure legends

**Supplementary Figure S1. Gene expression of *CjACOS5* genes in sugi.**

Relative mRNA levels of *CjACOS5a* and *CjACOS5b* in the investigated organs. Total RNA was isolated from stems (S), leaves (L), female strobili (FS), and male strobili (MS) in December. The average mRNA level of stems was defined as 1.0. The dots represent the relative mRNA level of each sample. The asterisks indicate significant differences between male strobili and the three other organs (Tukey–Kramer test; $n = 4$, $^{**}p < 0.01$).

**Supplementary Figure S2. PCR analysis of transgenic sugi trees.**

(a) PCR analysis of sugi cells (GE#1, GE#2, and GE#3) selected onto kanamycin-containing media. PCR products of *NPTII* were electrophoresed on an agarose gel and stained. WT is non-transgenic sugi cells. (b) PCR analysis of regenerated sugi plantlets. Genomic DNAs were prepared from leaves of non-transgenic sugi (WT) and transgenic sugi lines (GE). The arrows indicated the PCR products of *CjACOS5* (upper) and *NPTII* (lower).

**Supplementary Figure S3. Growth of *CjACOS5*-mutated transgenic sugi trees.**

(a) The investigated sugi trees were planted in June of 2017 and grown in a phytotron. (b–i) The photographs were taken in November of 2017. A non-transgenic sugi tree (b), GE#1-17 (c), GE#1-18 (d), GE#2-1 (e), GE#2-2 (f), GE#3-7 (g), GE#3-8 (h), and GE#3-9 (i). (j) The initial height was measured in November of 2017, at about 5 months after plantation. Plant height is shown for each group: WT (non-transgenic trees, $n = 6$), GE#1 (GE#1-17 and GE#1-18), GE#2 (GE#2-1–GE#2-6), and GE#3 (GE#3-7–GE#3-12). The dots represent the measured values of each tree. The asterisks indicate significant differences between the WT group and the transgenic tree groups (Dunnett's test; ***$p < 0.001$).

**Supplementary Figure S4. Mutations of *CjACOS5* genes in each strobilus.**

DNA sequence of PCR-amplified DNAs neighboring the break site of *CjACOS5* in a male strobilus which was sampled from the transgenic trees (GE#2-1, GE#2-2, and GE#3-7). The target sequence and PAM are shown in reversed black and underlined blue, respectively. The gaps (-) represent deleted nucleotides. Reads mean the number of sequenced DNAs after cloning of the PCR products.

**Supplementary Figure S5. Male strobili of genome-edited sugi trees in 2019.**

The sugi trees were grown in the phytotron (a–f) and in the special netted-house (g–i). The photo on the left is an outside (top) and a vertical section (bottom) of male strobili (scale bar, 1 mm). The photo on the right depicts pollen grains in razor-cutting suspension of male strobili (scale bar, 100 μm). Non-transgenic (WT) sugi strobili (a and g). Biallelic mutants of *CjACOS5*, GE#2-1 (b), GE#2-2 (c), GE#3-10 (f), GE#2-21 (h), and GE#2-22 (i). Chimeric mutants,

GE#3-7 (d) and GE#3-8 (e). The photographs were taken in February, 2019.

**Supplementary Figure S6. Male strobili of genome-edited sugi trees in 2020.**

The sugi trees were grown in the special netted-house. (a) Non-transgenic (WT) sugi male strobili (left; scale bar, 1 cm), magnified male strobili (middle; scale bar, 1 mm) and razor-cutting suspension of male strobili (right; scale bar, 100 μm). The magnified strobili are shown as an outside (top) and a vertical section (bottom). (b–e) Male strobili of *CjACOS5* biallelic mutant lines: GE#2-1 (b), GE#2-2 (c), GE#2-4 (d), and GE#2-6 (e). The photographs were taken in January, 2020.

**Supplementary Table S1. Synthetic DNAs used in this study.**

| Name of synthetic DNA | DNA sequence |
|---|---|
| For the cloning of the *CjACOS5* cDNAs using PCR | |
| CjACOS5_E1_F1 | 5′-ACACAAATTCATTTTCCCCATTGC-3′ |
| CjACOS5_ORF_R2 | 5′-CCTCATTAAAATTCTTATATTCTTTCATGCCATG-3′ |
| For the construction of the CRISPR/Cas9 vector | |
| CjACOS5_GE_FP | 5′-ATTGAAATGCTGTAAGAGAGGAAG-3′ |
| CjACOS5_GE_RP | 5′-AAACCTTCCTCTCTTACAGCATTT-3′ |
| For the PCR detection of *CjACOS5* | |
| CjACOS5_E1_F1 | 5′-ACACAAATTCATTTTCCCCATTGC-3′ |
| CjACOS5_E1_R1 | 5′-ATTCCGAGCACAATGATGGGATAC-3′ |
| For the PCR detection of *NPTII* | |
| NPT2U | 5′-GCTATTCGGCTATGACTGG-3′ |
| NPT2R | 5′-ATAGAAGGCGATGCGCTG-3′ |
| For RT-qPCR of *CjACOS5a* | |
| CjACOS5a_QPCR_LP1 | 5′-TCATTCTTCCAAATCTGGAGGTGAAG-3′ |
| CjACOS5a_QPCR_RP1 | 5′-TCTGTCTCTCGTGGCTTCTGATA-3′ |
| For RT-qPCR of *CjACOS5b* | |
| CjACOS5b_QPCR_LP1 | 5′-TCATTCTTCCAAATCTGGAGGTGAAA-3′ |
| CjACOS5b_QPCR_RP1 | 5′-CTGTCTCTCGTGGCTTCTGATG-3′ |
| For RT-qPCR of *CjeIF* | |
| CjeIF1_QPCR_LP1 | 5′-TACTGGTGGCCTTGCTGTATG-3′ |
| CjeIF1_QPCR_RP1 | 5′-GGAAACAGCGCAATGATCATTA-3′ |

**Supplementary Table S2. Mutation of *CjACOS5* in leaf DNA and pollen production in all transgenic sugi trees.**

| Line | DNA sequence of *CjACOS5a*[*1] | Mutation | Reads[*2] | DNA sequence of *CjACOS5b* | Mutation | Reads | Pollen[*3] |
|---|---|---|---|---|---|---|---|
| WT<br>(#13-8-2) | AAGTAAACGAAACTGAGCATGGC<br>GGACCAAAATGCTGTAAGAGAGG<br>AAGAGGAGATCA | None | ×7 | AAGGAAACGAAACTGAGCATGGC<br>GGACCAAAATGCTGTAAGAGAGG<br>AAGAGGAGATCA | None | ×4 | + |
| GE#1-17 | AAGTAAACGAAACTGAGCATGGC<br>GGACCAAAATGCTGTAAGAG---<br>AAGAGGAGATCA | 3-bp del | ×5 | AAGGAAACGAAACTGAGCATGGC<br>GGACCAAAATGCTGTAAGAGAG-<br>AAGAGGAGATCA | 1-bp del | ×5 | Withered |
| | AAGTAAACGAAACTGAGCATGGC<br>GGACCAAAATGCTGTAAGAGAG-<br>AAGAGGAGATCA | 1-bp del | ×6 | AAGGAAACGAAACTGAGCATGGC<br>GGACCAAAATGCTGTAAGAGAGG<br>-AGAGGAGATCA | 1-bp del | ×6 | |
| GE#1-18 | AAGTAAACGAAACTGAGCATGGC<br>GGACCAAAATGCTGTAAGAGAGG<br>AAGAGGAGATCA | None | ×16 | AAGGAAACGAAACTGAGCATGGC<br>GGACCAAAATGCTGTAAGAGAG-<br>AAGAGGAGATCA | 1-bp del | ×3 | Withered |
| | AAGTAAACGAAACTGAGCATGGC<br>GGACCAAAATGCTGTAAGAGAG-<br>AAGAGGAGATCA | 1-bp del | ×6 | | | | |
| GE#2-1 | AAGTAAACGAAACTGAGCATGGC | 1-bp del | ×7 | AAGGAAACGAAACTGAGCATGGC | 1-bp del | ×4 | − |

| | | | | | | | |
|---|---|---|---|---|---|---|---|
| | GGACCAAAATGCTGTAAGAGAG-<br>AAGAGGAGATCA | | | GGACCAAAATGCTGTAAGAGAG-<br>AAGAGGAGATCA | | | |
| GE#2-2 | AAGTAAACGAAACTGAGCATGGC<br>GGACCAAAATGCTGTAAGAGAG-<br>AAGAGGAGATCA | 1-bp del | ×3 | AAGGAAACGAAACTGAGCATGGC<br>GGACCAAAATGCTGTAAGAGAG-<br>AAGAGGAGATCA | 1-bp del | ×5 | – |
| GE#2-3 | AAGTAAACGAAACTGAGCATGGC<br>GGACCAAAATGCTGTAAGAGAG-<br>AAGAGGAGATCA | 1-bp del | ×5 | AAGGAAACGAAACTGAGCATGGC<br>GGACCAAAATGCTGTAAGAGAG-<br>AAGAGGAGATCA | 1-bp del | ×3 | – |
| GE#2-4 | AAGTAAACGAAACTGAGCATGGC<br>GGACCAAAATGCTGTAAGAGAG-<br>AAGAGGAGATCA | 1-bp del | ×4 | AAGGAAACGAAACTGAGCATGGC<br>GGACCAAAATGCTGTAAGAGAG-<br>AAGAGGAGATCA | 1-bp del | ×6 | – |
| GE#2-5 | AAGTAAACGAAACTGAGCATGGC<br>GGACCAAAATGCTGTAAGAGAG-<br>AAGAGGAGATCA | 1-bp del | ×4 | AAGGAAACGAAACTGAGCATGGC<br>GGACCAAAATGCTGTAAGAGAG-<br>AAGAGGAGATCA | 1-bp del | ×3 | – |
| GE#2-6 | AAGTAAACGAAACTGAGCATGGC<br>GGACCAAAATGCTGTAAGAGAG-<br>AAGAGGAGATCA | 1-bp del | ×4 | AAGGAAACGAAACTGAGCATGGC<br>GGACCAAAATGCTGTAAGAGAG-<br>AAGAGGAGATCA | 1-bp del | ×1 | – |

| | | | | | | | |
|---|---|---|---|---|---|---|---|
| GE#2-21 | AAGTAAACGAAACTGAGCATGGC GGACCAAAATGCTGTAAGAGAG-AAGAGGAGATCA | 1-bp del | ×2 | AAGGAAACGAAACTGAGCATGGC GGACCAAAATGCTGTAAGAGAG-AAGAGGAGATCA | 1-bp del | ×6 | − |
| GE#2-22 | AAGTAAACGAAACTGAGCATGGC GGACCAAAATGCTGTAAGAGAG-AAGAGGAGATCA | 1-bp del | ×2 | AAGGAAACGAAACTGAGCATGGC GGACCAAAATGCTGTAAGAGAG-AAGAGGAGATCA | 1-bp del | ×4 | − |
| GE#3-7 | AAGTAAACGAAACTGAGCATGGC GGACCAAAATGCTGTAAGAGAG-AAGAGGAGATCA | 1-bp del | ×11 | AAGGAAACGAAACTGAGCATGGC GGACCAAAATGCTGTAAGAGAGGAAGAGGAGATCA | None | ×4 | ± |
| | | | | AAGGAAACGAAACTGAGCATGGC GGACCAAAATGCTGTAAGAGAG-AAGAGGAGATCA | 1-bp del | ×1 | |
| GE#3-8 | AAGTAAACGAAACTGAGCATGGC GGACCAAAATGCTGTAAGAGAGGAAGAGGAGATCA | None | ×8 | AAGGAAACGAAACTGAGCATGGC GGACCAAAATGCTGTAAGAGAGGAAGAGGAGATCA | None | ×5 | + |
| | | | | AAGGAAACGAAACTGAGCATGGC GGACCAAAATGCTGTAAGAGAG-AAGAGGAGATCA | 1-bp del | ×4 | |

| | | | | | | | |
|---|---|---|---|---|---|---|---|
| GE#3-9 | AAGTAAACGAAACTGAGCATGGC GGACCAAAATGCTGTAAGAGAGG AAGAGGAGATCA | None | ×9 | AAGGAAACGAAACTGAGCATGGC GGACCAAAATGCTGTAAGAGAGG AAGAGGAGATCA | None | ×4 | ± |
| | AAGTAAACGAAACTGAGCATGGC GGACCAAAATGCTGTAAGAGAG- AAGAGGAGATCA | 1-bp del | ×1 | AAGGAAACGAAACTGAGCATGGC GGACCAAAATGCTGTAAGAGAG- AAGAGGAGATCA | 1-bp del | ×2 | |
|  | AAGTAAACGAAACTGAGCATGGC GGACCAAAATGCTGT-------- ----------------------- --------------TCTGGT | 45-bp del | ×1 |  |  |  |  |
| GE#3-10 | AAGTAAACGAAACTGAGCATGGC GGACCAAAATGCTGTAAGAGAG- AAGAGGAGATCA | 1-bp del | ×3 | AAGGAAACGAAACTGAGCATGGC GGACCAAAATGCTGTAAGAGAG- AAGAGGAGATCA | 1-bp del | ×7 | − |
| GE#3-11 | AAGTAAACGAAACTGAGCATGGC GGACCAAAATGCTGTAAGAGAG- AAGAGGAGATCA | 1-bp del | ×7 | AAGGAAACGAAACTGAGCATGGC GGACCAAAATGCTGTAAGAGAGG AAGAGGAGATCA | None | ×8 | ± |
| | | | | AAGGAAACGAAACTGAGCATGGC GGACCAAAATGCTGTAAGAGAG- AAGAGGAGATCA | 1-bp del | ×5 | |

| | | | | | | | |
|---|---|---|---|---|---|---|---|
| | | | | AAGGAAACGAAACTGAGCATGGCGGACCAAAATGCTGTAAGAGA---AGAGGAGATCA | 3-bp del | ×2 | |
| GE#3-12 | AAGTAAACGAAACTGAGCATGGCGGACCAAAATGCTGTAAGAGAG-AAGAGGAGATCA | 1-bp del | ×12 | AAGGAAACGAAACTGAGCATGGCGGACCAAAATGCTGTAAGAGAGGAAGAGGAGATCA | None | ×3 | ± |
| | | | | AAGGAAACGAAACTGAGCATGGCGGACCAAAATGCTGTAAGAGAG-AAGAGGAGATCA | 1-bp del | ×6 | |
| | | | | AAGGAAACGAAACTGAGCATGGCGGACatcaacagctatatattactcatGGAAGAGGAGATCA | 17-bp del and 24-bp ins | ×1 | |

[*1] The target sequence and PAM are shown in red and underlined blue, respectively. Gaps (-) are deleted nucleotides. Reversed small letters represent newly inserted nucleotides.

[*2] Reads mean the number of sequenced DNAs after cloning of the PCR products.

[*3] Pollen production is indicated as follows: +, strobili with pollen; −, strobili without pollen; ±, strobili with and without pollen; Withered, trees withered before flowering.

Supplementary Figure S1

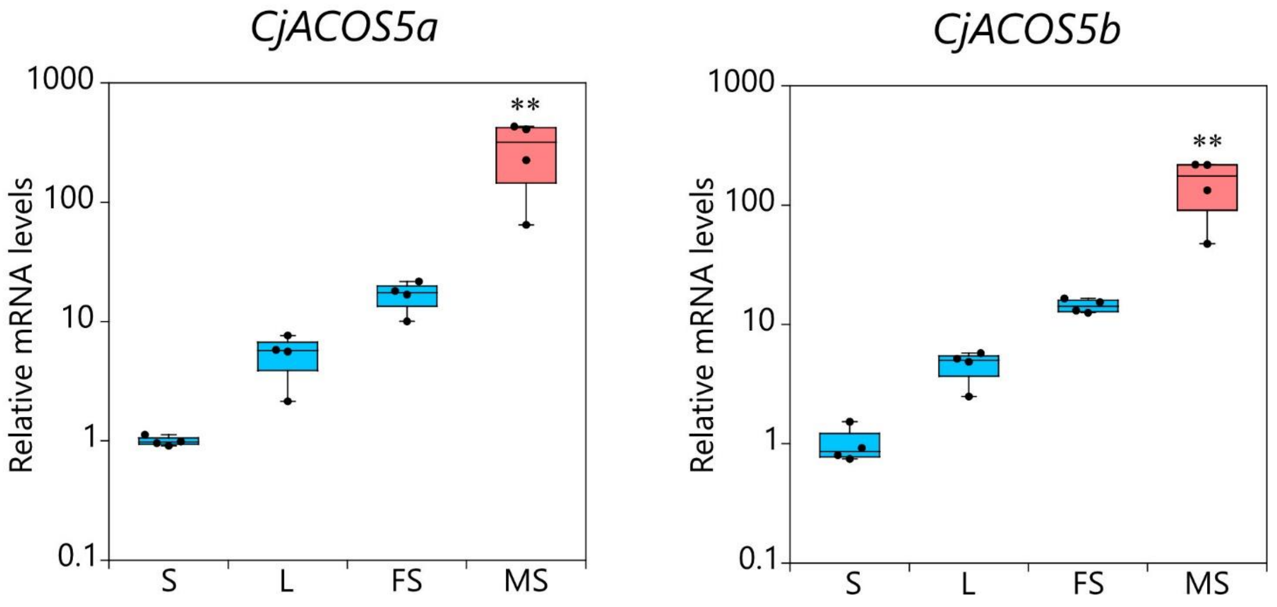

Supplementary Figure S2

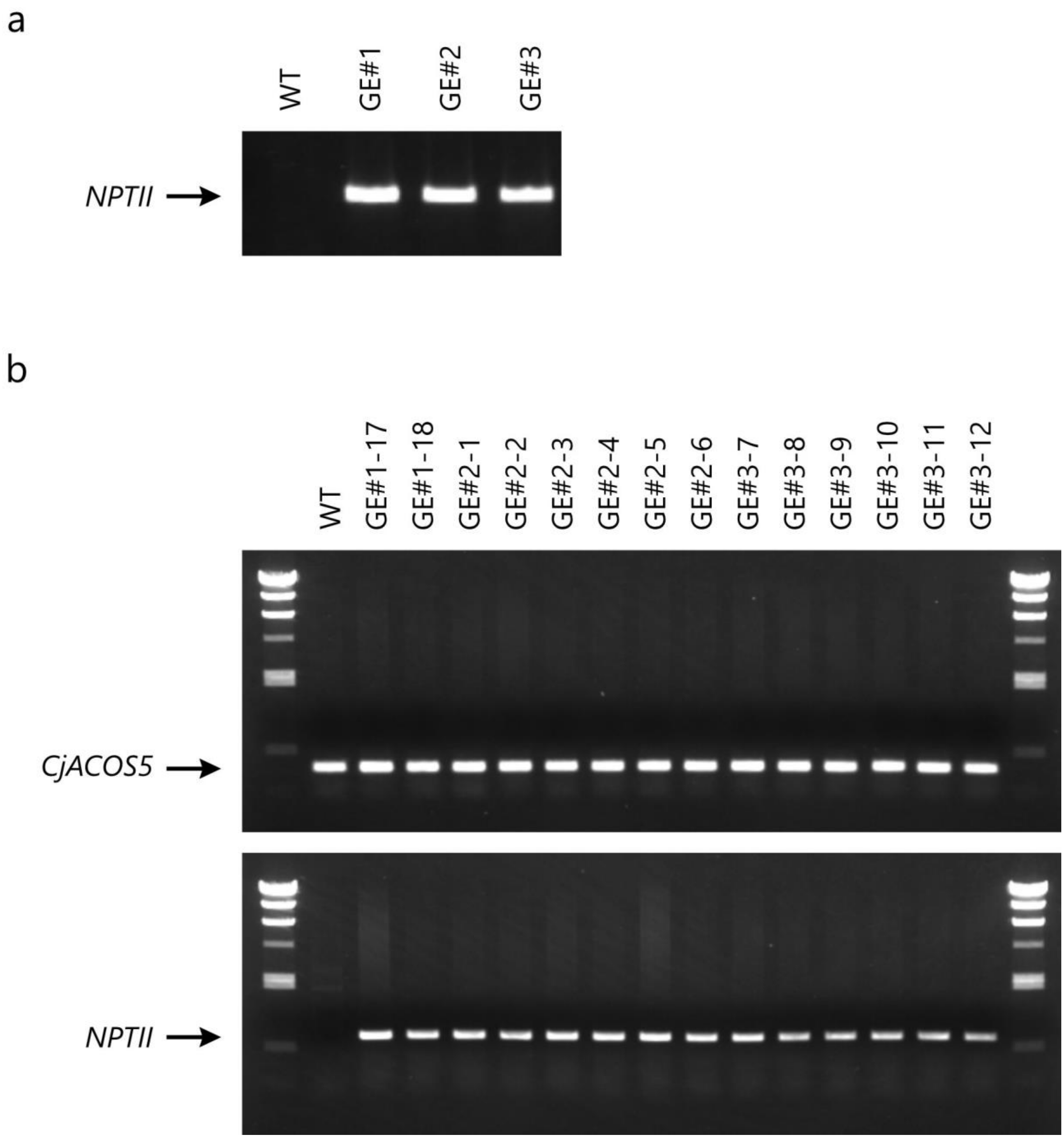

Supplementary Figure S3

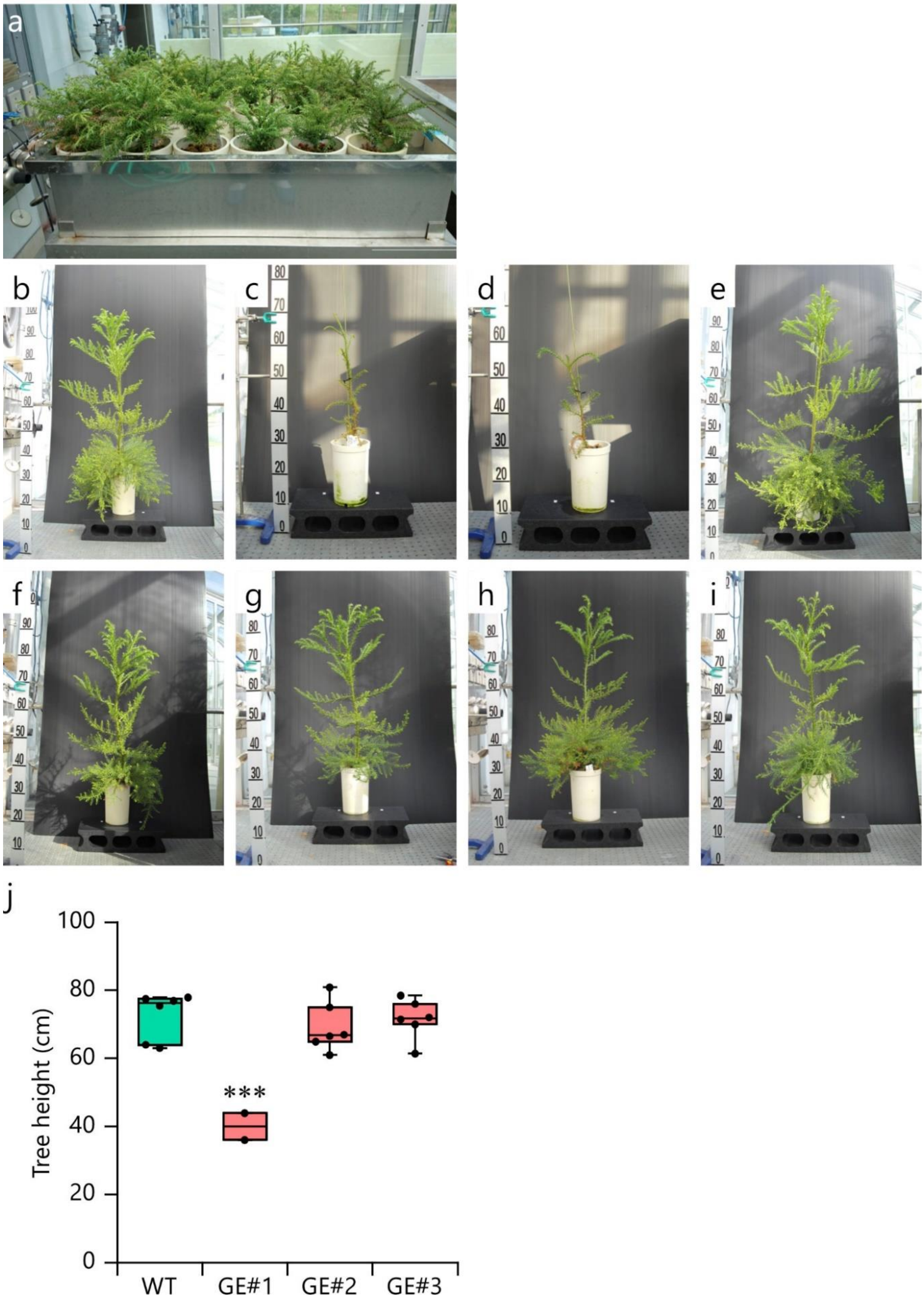

Supplementary Figure S4

| Samples | Genes | DNA Sequences | Reads |
|---|---|---|---|
| GE#2-1 Sterile male strobilus 1 | | | |
| | *CjACOS5a* | AAGTAAACGAAACTGAGCATGGCGGACCAAAATGCTGTAAGAGAG-AAGAGGAGATCA | ×6 |
| | *CjACOS5b* | AAGGAAACGAAACTGAGCATGGCGGACCAAAATGCTGTAAGAGAG-AAGAGGAGATCA | ×8 |
| GE#2-2 Sterile male strobilus 1 | | | |
| | *CjACOS5a* | AAGTAAACGAAACTGAGCATGGCGGACCAAAATGCTGTAAGAGAG-AAGAGGAGATCA | ×10 |
| | *CjACOS5b* | AAGGAAACGAAACTGAGCATGGCGGACCAAAATGCTGTAAGAGAG-AAGAGGAGATCA | ×11 |
| GE#3-7 Sterile male strobilus 1 | | | |
| | *CjACOS5a* | AAGTAAACGAAACTGAGCATGGCGGACCAAAATGCTGTAAGAGAGGAAGAGGAGATCA | ×1 |
| | | AAGTAAACGAAACTGAGCATGGCGGACCAAAATGCTGTAAGAGAG-AAGAGGAGATCA | ×7 |
| | *CjACOS5b* | AAGGAAACGAAACTGAGCATGGCGGACCAAAATGCTGTAAGAGAGGAAGAGGAGATCA | ×8 |
| | | AAGGAAACGAAACTGAGCATGGCGGACCAAAATGCTGTAAGAGA--AAGAGGAGATCA | ×1 |
|  |  | AAGGAAACGAAACTGAGCATGGCGGACCAAAATGCTGTAAGA----AAGAGGAGATCA | ×1 |
| GE#3-7 Sterile male strobilus 2 | | | |
| | *CjACOS5a* | AAGTAAACGAAACTGAGCATGGCGGACCAAAATGCTGTAAGAGAG-AAGAGGAGATCA | ×5 |
| | *CjACOS5b* | AAGGAAACGAAACTGAGCATGGCGGACCAAAATGCTGTAAGAGAGGAAGAGGAGATCA | ×11 |
| | | AAGGAAACGAAACTGAGCATGGCGGACCAAAATGCTGTAAGAGAG-AAGAGGAGATCA | ×1 |
| GE#3-7 Fertile male strobilus 1 | | | |
| | *CjACOS5a* | AAGTAAACGAAACTGAGCATGGCGGACCAAAATGCTGTAAGAGAGGAAGAGGAGATCA | ×1 |
| | | AAGTAAACGAAACTGAGCATGGCGGACCAAAATGCTGTAAGAGAG-AAGAGGAGATCA | ×7 |
| | *CjACOS5b* | AAGGAAACGAAACTGAGCATGGCGGACCAAAATGCTGTAAGAGAGGAAGAGGAGATCA | ×7 |
| | | AAGGAAACGAAACTGAGCATGGCGGACCAAAATGCTGTAAGAGAG-AAGAGGAGATCA | ×3 |
| GE#3-7 Fertile male strobilus 2 | | | |
| | *CjACOS5a* | AAGTAAACGAAACTGAGCATGGCGGACCAAAATGCTGTAAGAGAG-AAGAGGAGATCA | ×6 |
| | *CjACOS5b* | AAGGAAACGAAACTGAGCATGGCGGACCAAAATGCTGTAAGAGAGGAAGAGGAGATCA | ×10 |

Supplementary Figure S5

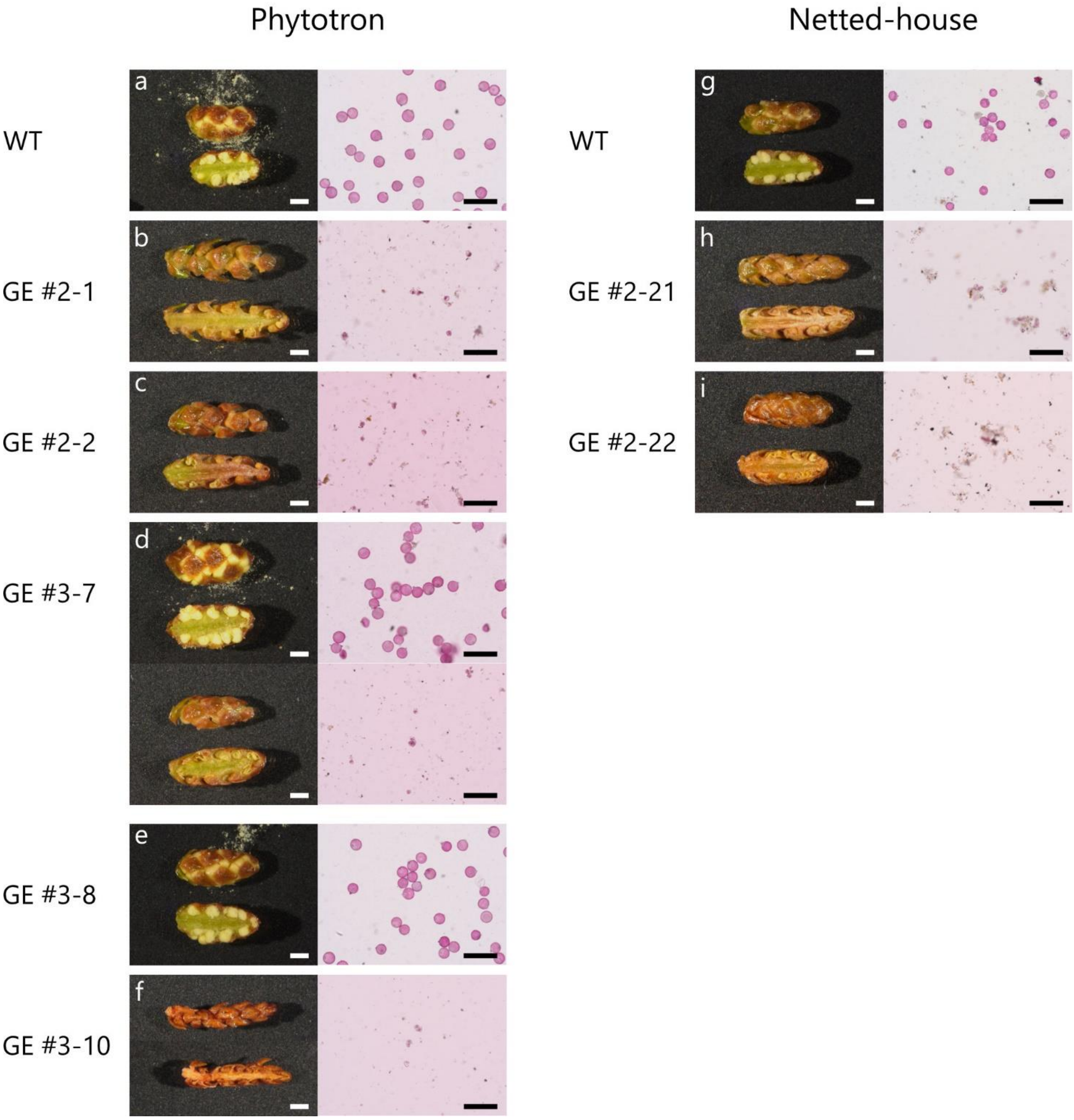

Supplementary Figure S6

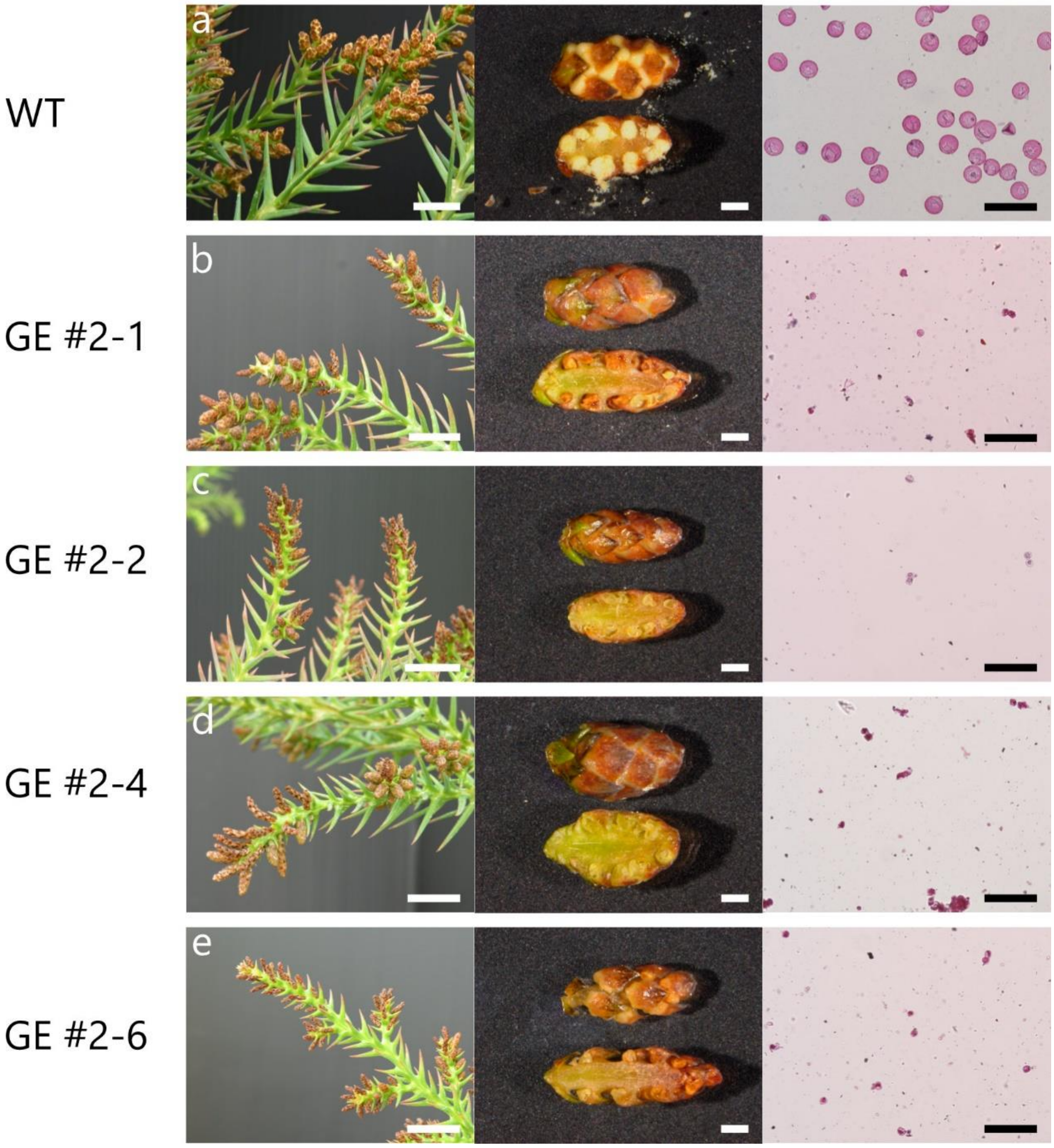
